# Introducing prescribed biases in out of equilibrium Markov models

**DOI:** 10.1101/198697

**Authors:** Purushottam D. Dixit

## Abstract

Markov models are often used in modeling complex out of equilibrium chemical and biochemical systems. However, many times their predictions do not agree with experiments. We need a systematic framework to update Markov models to make them consistent with constraints that are derived from experiments. Here, we present a framework based on the principle of maximum path entropy to update Markov models using stationary state and dynamical trajectory-based constraints. We illustrate the framework using a biochemical model network of growth factors-based signaling. We also show how to find the closest detailed balanced Markov model to a given Markov model. Further applications and generalizations are discussed.

## Introduction

Many chemical and biochemical phenomena of interest are driven by external forces and operate away from thermodynamic equilibrium. Examples include cellular metabolism^(1)^, biochemical signaling networks^(2)^, and elemental cycles in the atmosphere^(3)^.

Markov models are ubiquitously used in modeling such non-equilibrium processes. Quite often, ‘prior’ Markov models do not agree with experimental data. At the same time, the data is coarse-grained and cannot uniquely ‘update’ models to make them consistent with data. Here is an example. Consider an out of equilibrium network of biochemical reactions. The network is modeled using a chemical master equation (CME). Using the CME, we have predicted the steady state abundances of individual species in the network. Experiments often measure averages or distributions of abundaces of network species as well. Now imagine that the coarse grained measurements do not agree with the model predictions. How do we uniquely modify the master equation to reproduce the measured abundances? Indeed, for this problem and for many others, a conceptual framework that allows us to systematically incorporate user specified constraints and update out of equilibrium Markov models is needed.

The inference principle of maximum entropy (ME) is ideal to model and update probability distributions with incomplete information ^(4,5)^. ME, introduced in statistical physics nearly a century ago, is now widely used across multiple areas of physics, chemistry, and biology. Most popular applications of ME predict probabilities of states of a system from incomplete information. The dynamical version of ME (maximum *path* entropy or maximum caliber) can be used to estimate probabilities over trajectories. Previously, we have used the maximum path entropy framework to study Markovian systems^(6–8)^. We derived the transition rates of Markov models constrained to reproduce a stationary distribution and a few dynamical observables^(6,7)^. We showed that the predicted rates can accurately capture the kinetics of transitions among metastable states of complex biomolecular systems. We have also derived both the stationary distribution as well as the transition rates of a Markov model that was constrained to reproduce only dynamical observables. Notably, we showed that state space topology has a significant impact on the stationary distribution^(8)^.

In this work, we present a ME based framework to update prior non-equilibrium Markov models subject to state and dynamical trajectory-based constraints. Then we illustrate the framework using a biochemical network that represents growth factor based signaling. We conclude with possible future directions.

## Updating Markov models

We consider a discrete time discrete state Markovian system. We denote by {*a*} the states of the system and by {*q*_*ab*_} the transition probabilities among the states. We assume that *q*_*ab*_ > 0 if the transition *a* → *b* is allowed in a single time step and zero otherwise. Finally, we assume that the Markov model is aperiodic and irreducible.

Dynamical observables *r*_*ab*_ can be defined over the transitions *a* → *b* of the trajectories generated by the Markov process^(6–8)^. Examples of *r*_*ab*_ include total distance traversed by an agent on a graph, the number of spin changes in a unit time step of an Ising model, the number of amino acid structural contacts broken by a protein in a single time step, etc. The ensemble average of *r*_*ab*_ over stationary state paths of the Markov process is given by

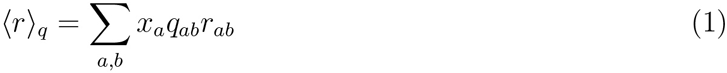

where {*x*_*a*_} is the stationary distribution of the Markov model. The subscript *q* denotes that the average was taken over a Markov model with transition probabilities {*q*_*ab*_}. The stationary distribution {*x*_*a*_} satisfies

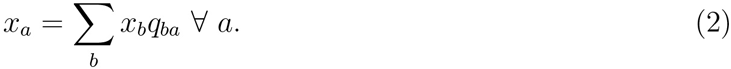

We note that *r*_*ab*_ includes purely state-dependent observables as well. For example, the ensemble average of a state-dependent property *f*_*a*_ is expressed as

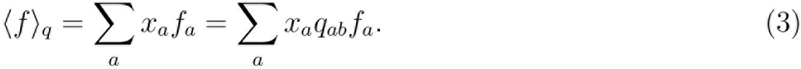

Consider that the experimental measurement *r̅* of the average of *r*_*ab*_ does not agree with the model prediction ⟨*r*⟩_*q*_. How do we fix/update the Markov model so that the two averages agree? We seek a *least perturbed* Markov model relative to the prior model. We take the maximum relative path entropy (minimum Kullback-Leibler divergence) approach. The relative path entropy is given by^(9)^

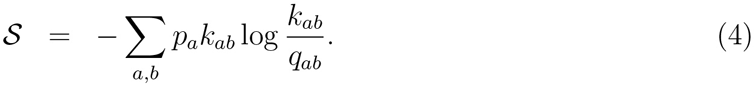

In Eq. 4, {*p*_*a*_} is the updated stationary distribution and {*k*_*ab*_} are the updated transition probabilities. We maximize *S* in Eq. 4 subject to following constraints,

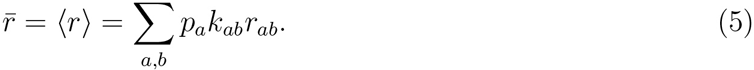

Additionally, {*p*_*a*_} and {*k*_*ab*_} are not mutually independent. We have

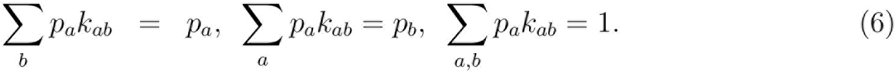

The three types of constraints in Eq. 6 reflect probability normalization, stationarity of {*p*_*a*_} with respect to the transition probabilities {*k_ab_*}, and normalization of the stationary distribution {*p*_*a*_} respectively.

We carry out the maximization using the method of Lagrange multipliers. We write the Caliber^(8)^

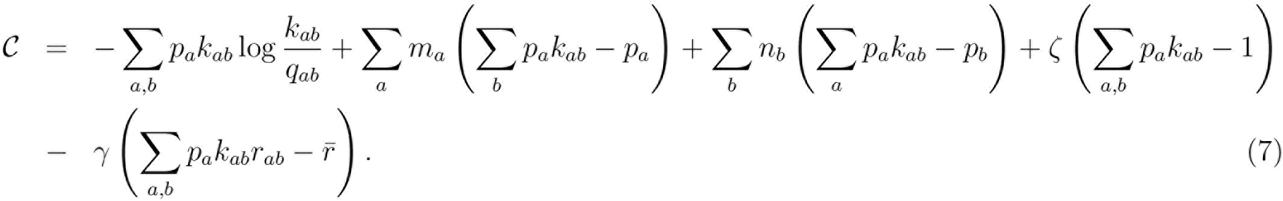

Differentiating *C* with respect to *k*_*ab*_ and setting the derivative to zero,

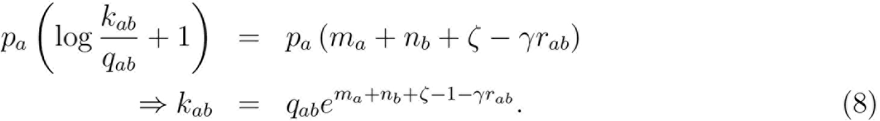

Differentiating *C* with respect to *p*_*a*_ and setting the derivative to zero,

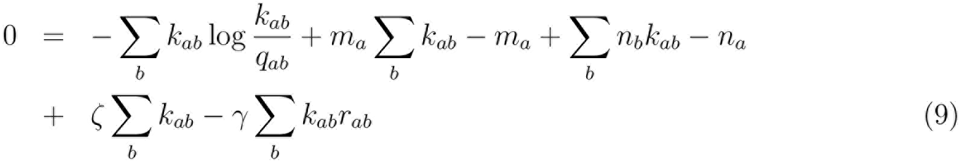

Substituting *k*_*ab*_ from Eq. 8 and using Σ_*b*_ *k*_*ab*_ = 1, we get

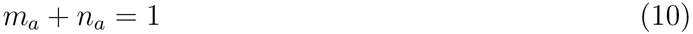

and consequently,

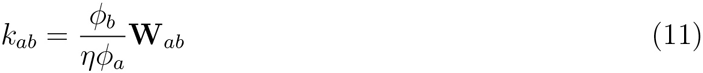

where

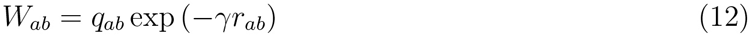

In Eq. 11, *ϕ*_*a*_ = *e*^−*m*_*a*_^ and *η* = *e*^-*ζ*^ are the modified Lagrange multipliers. We determine *ϕ*_*a*_ by imposing Σ_*b*_ *k*_*ab*_ = 1. We have,

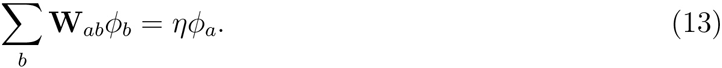

In other words, 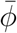 is the eigenvector of matrix **W** with eigenvalue *η*. There are *n* choices for the eigenvalue and the corresponding eigenvector. To guarantee positivity of the transition probabilities *k_ab_* in Eq. 11, we choose *η* to be the Perron-Frobenius eigenvalue. Note that the existence of the Perron-Frobenius eigenvalue is guaranteed because the Markov process is assumed to be aperiodic and irreducible. The eigenvector 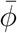 corresponding to the Perron-Frobenius eigenvalue *η* is such that *ϕ*_*a*_ > 0 ∀ *a*.
Next, we determine the stationary distribution {*p*_*a*_}. We have

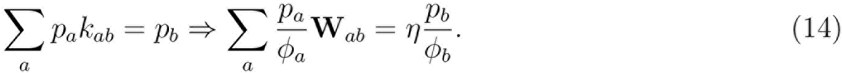

We write *ψ*_*a*_ = *P*_*a*_/*ϕ*_*a*_,

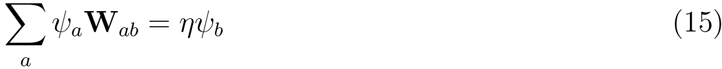

In other words, if 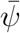 is the left Perron-Frobenius eigenvector and 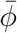 is the right Perron-Frobenius eigenvector of **W** with the same eigenvalue *η*, the stationary distribution is given by the outer product

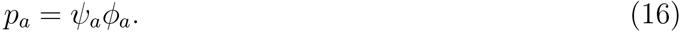

Eq. 11 and Eq. 16 complete our derivation. To summarize, we start from a prior Markov model described by transition probabilities {*q*_*ab*_} and a stationary distribution {*x*_*a*_}. Based on a dynamical constraint *r̅*, we update it to a Markov model described by transition probabilities {*k*_*ab*_} and a stationary distribution {*p*_*a*_}. In practice, we construct Markov models for multiple values of γ, the Lagrange multiplier that sets the value of the dynamical constraint ⟨*r*⟩ = *r̅*. For a given γ, we construct the matrix **W** (see Eq. 12). Next we find its Perron-Frobenius eigenvalue *η* and the corresponding eigenvector 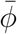. Next we calculate the transition probabilities {*k*_*ab*_} (see Eq. 11). We vary 7 until the dynamical constraint ⟨*r*⟩ = *r̅* is satisfied. The updated model is consistent with the imposed constraint and has the maximum relative entropy with respect to the prior model. A generalization to multiple constraints 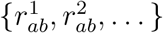 is straightforward. We introduce one Lagrange multiplier γ_*i*_ for each constraint 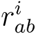. We have

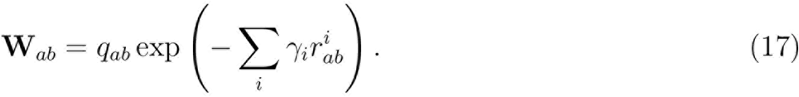

The rest of the procedure is similar to the one described in the paragraph above.

In the next two sections, we illustrate our method with a couple of examples.

## Illustrative example 1: Growth factor activated receptor tyrosine kinase (RTK) cascade

Growth factors are secreted peptides that bind to growth factor receptors on plasma membranes of cells. Upon binding with the growth factor ligand, the receptors get phosphorylated (activated) and transmit the phosphorylation signal downstream. Ligand-free (empty) receptors are delivered to the cell surface from the cytoplasm. Empty, bound, and active state receptor on the plasma membrane can be internalized into the cytosol and degraded. A key feature of many growth factor activated pathways is preferential internalization of active receptors^(10)^. Signaling cascades downstream of growth factor receptors are responsible for a wide array of phenotypes in humans including upregulation of metabolism, proliferation, and cell motility^(11)^.

At steady state, the total number of growth factor receptors on the plasma membrane in the absence of extracellular ligand ranges between *N*_*r*_ ~ 10^4^ – 10^6^. To model interactions of the growth factor receptor with the ligand, we imagine a ‘site’ for each receptor that can take four states. State ‘1’ represents the site without a receptor, state ‘2’ represents the site with an empty receptor, state ‘3’ represents ligand bound receptor, and state ‘4’ represents ligand bound ‘active’ receptor. Fig. 1 shows a schematic of the biochemical network. The arrows represent all possible single-step transitions.

**Figure 1:**
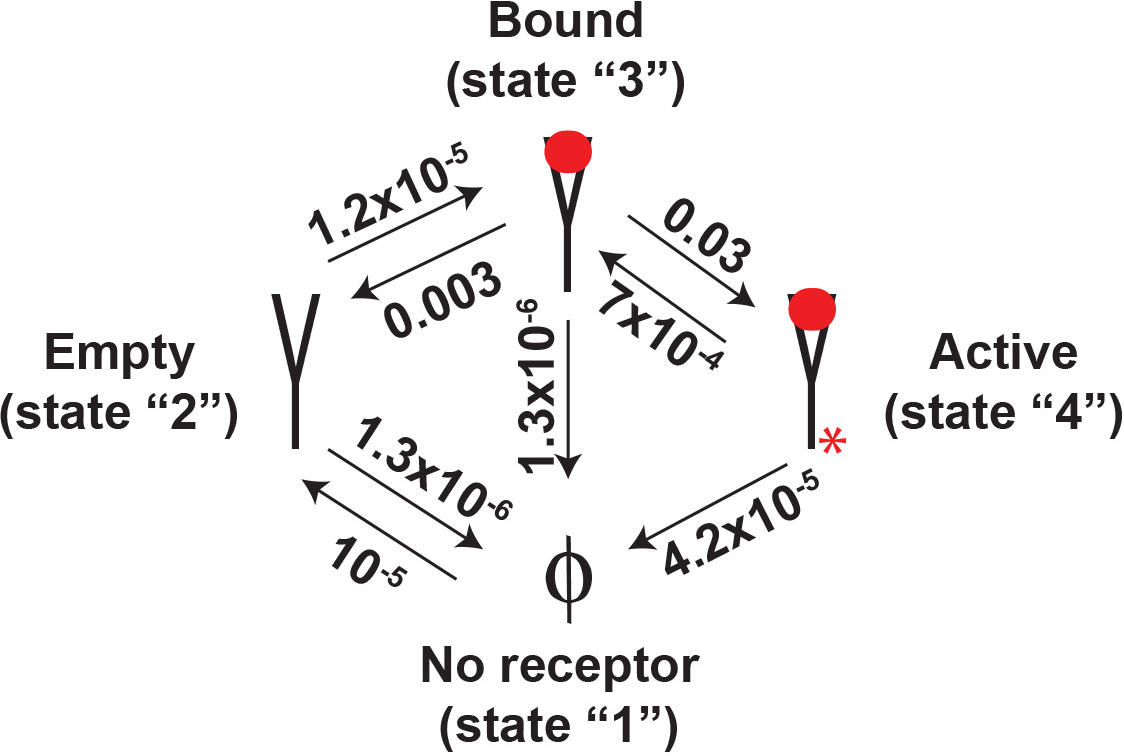
A four state model of the growth factor activated pathway. Empty receptors on the plasma membrane can be bound by a ligand and then phosphorylated (active). Empty, ligand-bound, as well as active receptors are internalized and degraded. The internalization rate of active receptors is faster than that of empty and bound receptors. Finally, empty receptors can be delivered to the plasma membrane from the cytoplasm. The transition probabilities are shown next to the arrows, we have assumed [*L*] = 1 ng/ml.

We model the transition probabilities using the experimentally measured microscopic rates^(10,12,13)^

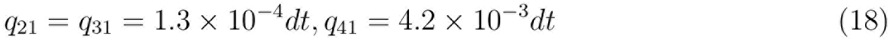

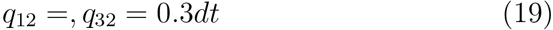

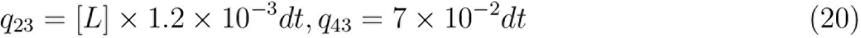

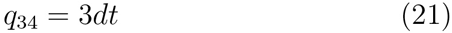

where *dt* = 0.01 seconds. [*L*] is the extracellular ligand concentration measured in ng/ml. We assume that [*L*] is held constant at [*L*] = 1 ng/ml. We also assume that the total number of cell surface receptors in the absence of any ligand is *N*_r_ = 4 × 10^5^ per cell^(14)^. Note that the system described by Eqs. 21 is inherently out of equilibrium; empty receptors are constantly shuttled to the plasma membrane. The empty receptors are eventually converted into active receptors. Active receptors are internalized and degraded at a faster rate.

An important quantity in RTK networks is the total number of active receptors *N*_act_ = *p*_4_*N*_r_ at steady state. For the transition probabilities given in Eq. 21, at steady state, we have *N*_act_ = 0.0618 × *N*_r_ ≈ 2.5 × 10^4^. Notably, RTK networks are frequently mutated in cancers. Moreover, mutations often lead to highly active networks^(11)^. Imagine a situation where the experiments suggest that the number of active receptors at steady state is *N*_act_ = 10^5^ (or equivalently *p*_4_ = *N*_act_/*N*_r_ = 0.25). How do we update the transition rates {*q*_*ab*_} to reflect this change?

The constraint *p*_4_ = 0.25 can be represented as ⟨*r*⟩_*k*_ = Σ*p*_*a*_*k*_*ab*_*r*_*ab*_ where

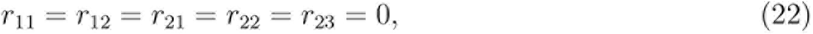

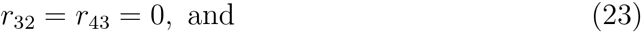

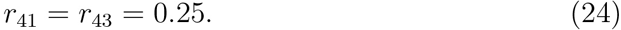

We constructed multiple Markov models by incorporating the constraint ⟨*r*⟩_*k*_ at multiple values of the Lagrange multiplier γ used to impose the constraint. To do so, we first constructed the matrix **W** (see Eq. 12) at each value of γ. Next, we determined the Perron-Frobenius eigenvalue *η* and the corresponding eigenvector 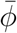 and 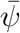. Finally, we calculated the updated transition probabilities {*k*_*ab*_} (see Eq. 11). In Fig. 2, we plot the relative log ratio log *k*_*ab*_/*q*_*ab*_ of the updated transition probabilities *k*_*ab*_ and the corresponding prior probabilities *q*_*ab*_ as a function of γ. Notably, the constraint in Eq. 24 changes all transition probabilities in a non-trivial manner. Moreover, while some some transition probabilities are insensitve to variation in the Lagrange multiplier (see Fig. 2f,g) others can vary over several orders of magnitude (see Fig. 2e,h).

**Figure 2:**
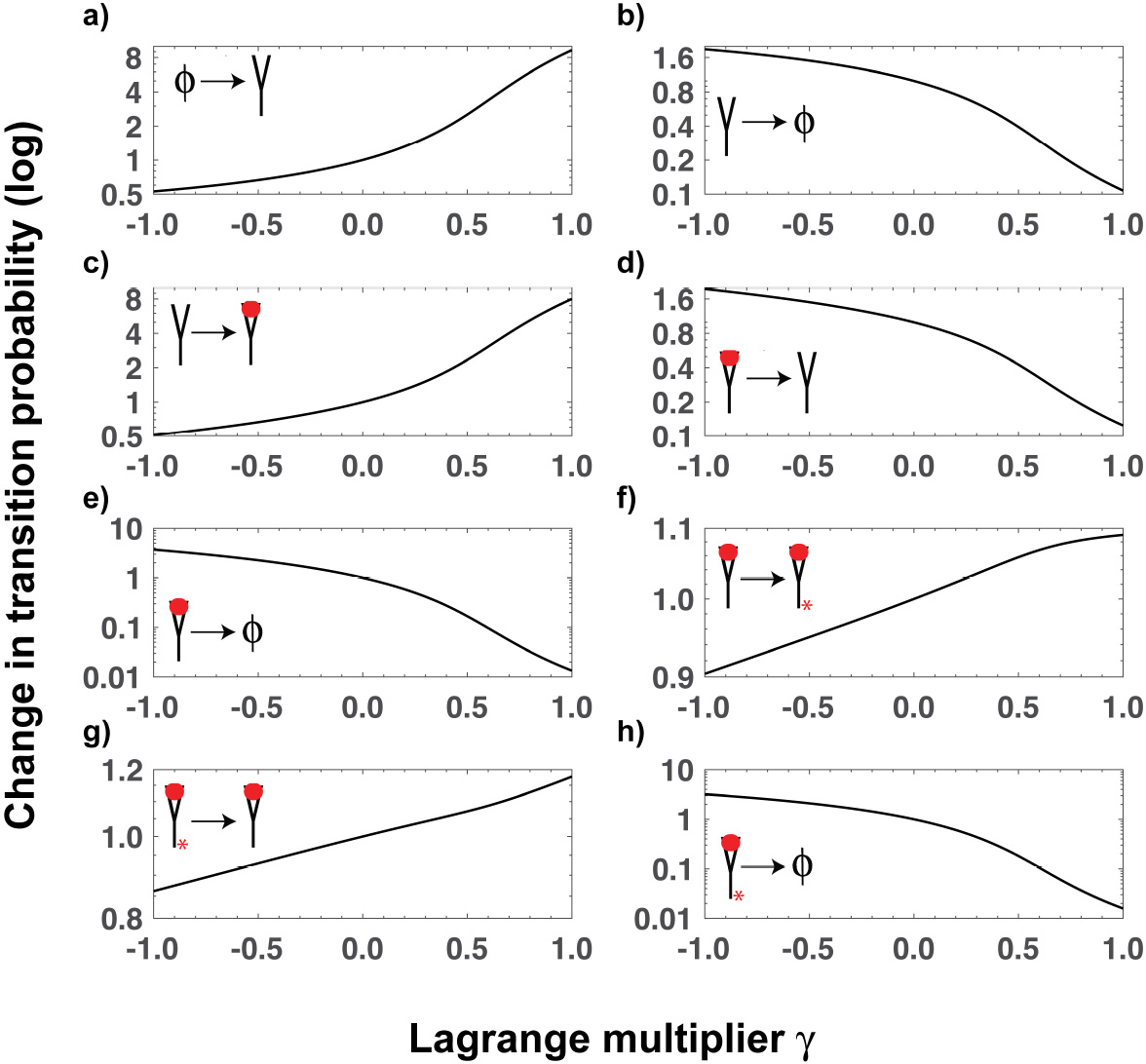
The ratio *k*_*ab*_/*q*_*ab*_ of the updated transition probabilities *k*_*ab*_ to the prior transition probabilities *q*_*ab*_ as a function of the Lagrange multiplier γ.

For each of the Markov model (corresponding to a fixed Lagrange multiplier γ) we evaluate the stationary distribution *P*_*a*_ = *ϕ*_*a*_ψ_*a*_ (see Eq. 16). In Fig. 3, we plot the predicted *p*_4_(γ) as a function of the Lagrange multiplier γ. As expected, *P*_4_(γ = 0) = 0.0618. We find that at *p*_4_(γ = γ* = 0.4) = 0.25. In Table 1 we tabulate the modified transition probabilities {*k*_*ab*_} for γ = γ*. Notably, the updated model predicts that the most perturbed transition probabilities are those of internalization of ligand-free (2 → 1), ligand-bound (3 → 1), and active receptors (4 → 1). Indeed, receptor mutations that lead to lowered receptor internalization rates are a well documented reason of sustained RTK activity in many type of cancers^(15)^. These predictions can be tested experimentally by measuring the microscopic rates of internalization^(10)^.

**Figure 3:**
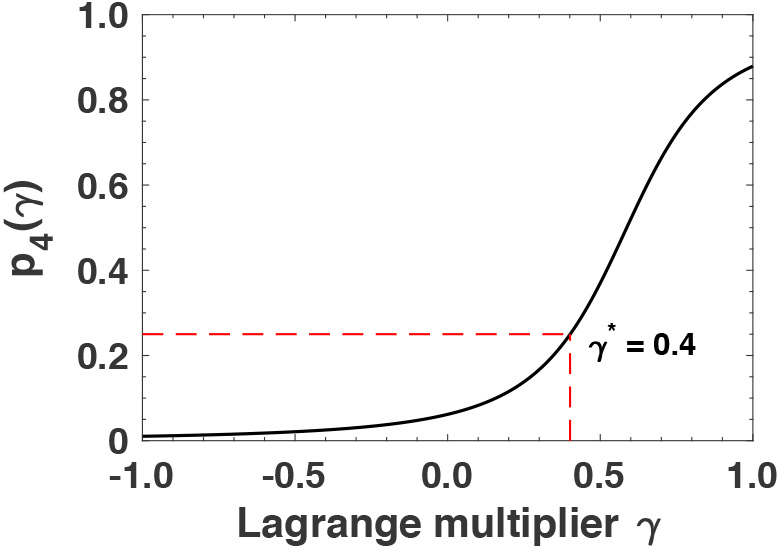
The steady state probability of active receptors *P*_4_(γ) as a function of the Lagrange multiplier γ. The dashed red lines show the value of γ ~ 0.4 corresponding to *p*_4_(γ) = 0.25.

**Table 1:** A comparison between the prior transition probabilities *q*_*ab*_ and the updated transition probabilities *k*_*ab*_. The updated probabilities were obtained by setting *p*_4_ = 0.25.

| Transition $a \rightarrow b$ | prior $q_{ab}$ | updated $k_{ab}$ | Ratio $k_{ab}/q_{ab}$ |
| --- | --- | --- | --- |
| $1 \rightarrow 2$ | $10^{-5}$ | $1.9 \times 10^{-5}$ | 0.9 |
| $2 \rightarrow 1$ | $1.3 \times 10^{-6}$ | $6.6 \times 10^{-7}$ | 0.5 |
| $2 \rightarrow 3$ | $1.2 \times 10^{-5}$ | $2.2 \times 10^{-5}$ | 1.8 |
| $3 \rightarrow 2$ | $3 \times 10^{-3}$ | $1.6 \times 10^{-3}$ | 0.5 |
| $3 \rightarrow 1$ | $1.3 \times 10^{-6}$ | $3.6 \times 10^{-7}$ | 0.3 |
| $3 \rightarrow 4$ | $3 \times 10^{-2}$ | $3.14 \times 10^{-2}$ | 1.0 |
| $4 \rightarrow 3$ | $7 \times 10^{-4}$ | $7.4 \times 10^{-4}$ | 1.0 |
| $4 \rightarrow 1$ | $4.2 \times 10^{-5}$ | $1.2 \times 10^{-5}$ | 0.3 |

## Illustrative example 2: Finding the ‘nearest’ detailed balanced Markov model

In this section, we show how to construct a detailed balanced Markov model that is least perturbed relative to a prior out of equilibrium Markov model. One motivation is as follows. Consider a system with discrete states {*a*} at thermodynamic equilibrium with its surroundings. Imagine that you have estimated a Markov model from time series data on this system. Let the transition probabilities of the model be {*q*_*ab*_} and the stationary distribution be {*x*_*a*_}. A common example is the popular Markov state modeling (MSM) approach to study biomolecular dynamics^(16)^. Often, the molecular dynamics trajectories are not completely converged. This leads to erroneous estimation of transition probabilities as well as the stationary distribution. In this case, it is likely that the empirically estimated stationary distribution does not satisfy detailed balance with respect to the empirically estimated transition probabilities, i.e.

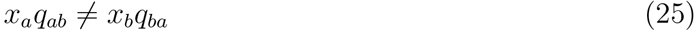

at least for some *a* and *b*. How do we fix the empirical Markov model so that it is detailed balanced? Here too, we find a solution within the maximum relative entropy framework. We maximize the relative entropy

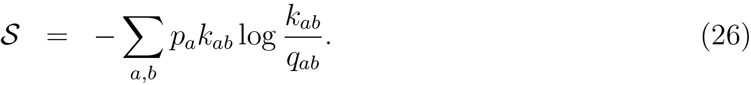

subject to constraints in Eq. 6 and

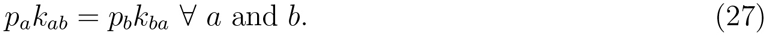

The last set of constraints imposes detailed balance in the updated model. We write the Caliber,

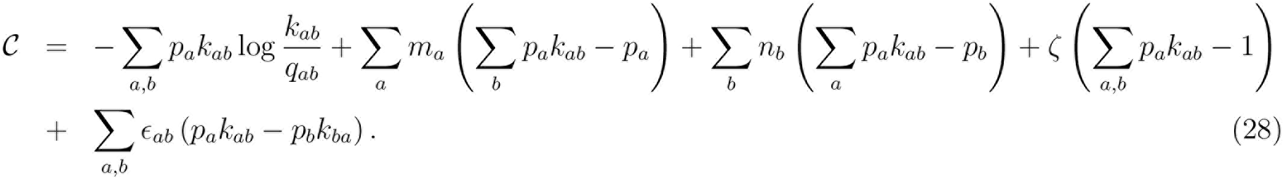

In Eq. 28, the Lagrange multipliers {*ϵ*_*ab*_} impose detailed balance between states *a* and *b*.

We maximize the caliber with respect to the transition probabilities {*k*_*ab*_} and the stationary distribution {*p*_*a*_} (see appendix for details). As above (see Eq. 11), we have

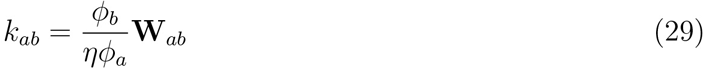

where

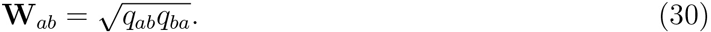

Note that **W** is symmetric in *a* and *b*. *η* is the Perron-Frobenius eigenvalue of **W** and 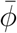 is the corresponding eigenvector. The stationary distribution is given by

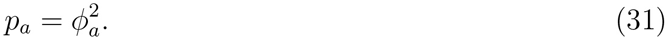

It is easy to check that {*p*_*a*_} satisfies detailed balance with respect to {*k*_*ab*_}. We have

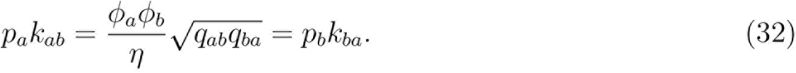

Eqs. 29, 30, and 31 complete the derivation of detailed balanced Markov model that has the least Kullback-Leibler divergence compared to a prior Markov model. Notably, the updated Markov model changes both the transition probabilities as well as the stationary distribution. We leave it for future studies to investigate whether the above procedure leads to better predictions of free energy landscapes and kinetics from simulated data.

## Possible generalizations

We envision that a major application of this work is in modeling non-equilibrium molecular machines^(17)^ as well as stochastic chemical reaction networks that are described using the chemical master equation (CME)^(18)^. Notably, in both these cases, various transition probabilities are not independent of each other but are rather parametrized from a smaller set of rates. For concreteness, consider the simple CME describing stochastic evolution of the probability *p*(*n*; *t*) of observing *n* mRNA transcripts^(19)^. We have

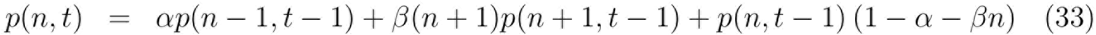

In Eq. 33, *α* is the probability of transcription and *β* is the probability of degradation of the transcript. In Eq. 33, the transitions *n* – 1 → *n* and *n* → *n* + 1 involve the same process (mRNA transcription) and are described by the same rate *α*. Similar is the case for transitions involving transcript degradation. If the framework described here is employed naively to update the Markov model described by Eq. 33, the updated model is not guaranteed to have these constitutive relationships. However, a straightforward generalization will allow us to handle parametrized Markov models. Here, instead of maximizing the relative entropy with respect to the full transition probability matrix {*k*_*ab*_}, we instead recognize that the transition probabilities {*k*_*ab*_} are parametrized and carry out the entropy maximization with respect to those parameters. For example, in case of Eq. 33, we will find updated parameters *α* → *α′* and *β* → *β′* such that the updated CME has the maximum relative entropy with respect to the prior CME and satisfies imposed constraints.

## Discussions

Many complex chemical and biological systems operate out of equilibrium. There are two general issues with modeling their dynamics. First, given the complexity of the dynamics, first principles models often do not agree with experimental data. Second, the experimental data are often coarse grained; measurements consists of averages of a few variables. As a result, one cannot uniquely update prior Markov models to make them consistent with data. In this work, we presented a framework based on the principle of maximum relative entropy (minimum Kullback-Leibler divergence) to uniquely update Markove models based on constraints. We used the framework to update the transition probabilities in a model of a receptor tyrosine kinase (RTK). We also showed how to obtain the closest detailed balanced Markov model to a given Markov model. We envision that this framework will be useful in modeling out of equilibrium chemical and biochemical networks.

## Appendix Imposing detailed balance

We start with Eq. 28,

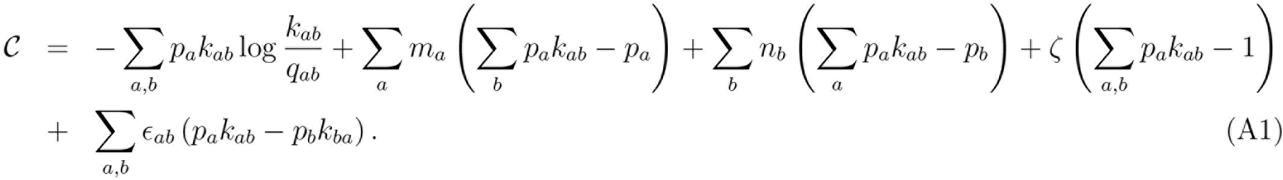

We have introduced Lagrange multipliers *ϵ*_*ab*_ to enforce detailed balance. As above, all summations involving two indices are restricted to edges of the graph.

Differentiating the Caliber with respect to *k*_*ab*_, we have

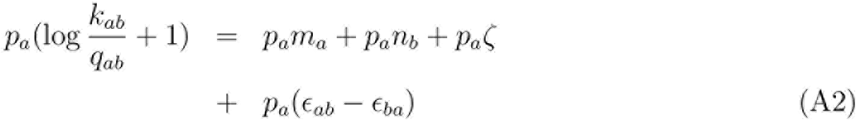

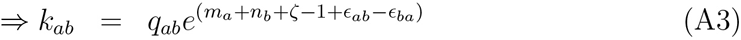

Differentiating the Caliber with respect to *p*_*a*_, we have

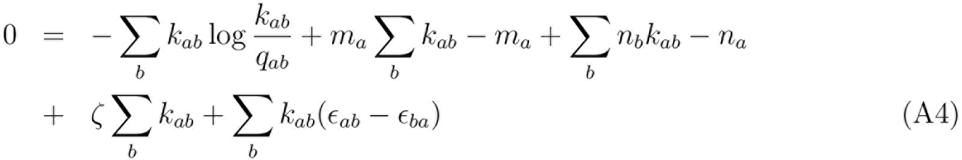

Substituting *k*_*ab*_ from Eq. A3, we get

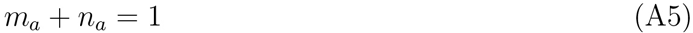

Substituting in Eq. A3, we get

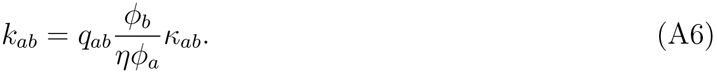

Here, *ϕ*_*a*_ = *e*^*-m*_*a*_^, *η* = *e*^*-ζ*^, and *K*_*ab*_ = *e*^*ϵ*_*ab*_–*ϵ*_*ba*_^. Notice that *K*_*ab*_*K*_*ba*_ = 1.
o determine *K*_*ab*_, we impose detailed balance,

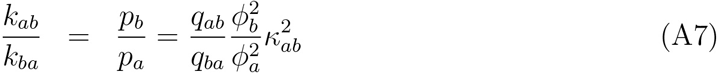

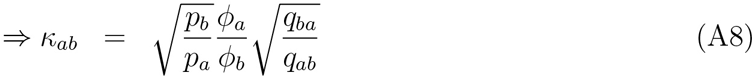

Tus, the transition probabilities are

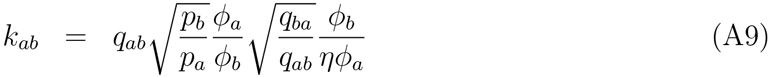

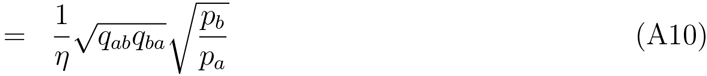

Let 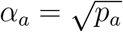 and 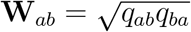. Using Σ_*b*_ *k*_*ab*_ =1, we have

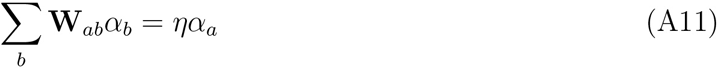

Thus, *ᾱ*, the vector of square roots of probabilities is the eigenvector of **W** with eigenvalue *η*.

